# Functional interrogation of a depression-related serotonergic SNP, rs6295, using a humanized mouse model

**DOI:** 10.1101/472621

**Authors:** Ashley M. Cunningham, Tabia L. Santos, Vanessa A. Gutzeit, Heather Hamilton, René Hen, Zoe R. Donaldson

## Abstract

The serotonin 1A receptor (5-HT1A) system has been extensively implicated in modulating mood and behavior. Notably, 5-HT1A levels in humans display remarkable variation and differences in receptor levels have been linked with a variety of psychiatric disorders. Further, manipulation of receptor levels in mice suggests that changes in receptor levels that model existing human variation are sufficient to drive behavioral alterations. As a result, genetic mechanisms that modulate human 5-HT1A levels may be important for explaining individual differences in mood and behavior, representing a potential source of psychiatric disease risk. One common genetic variant implicated in differential 5-HT1A levels is the G/C single nucleotide polymorphism (SNP), rs6295, located upstream of the human 5-HT1A gene. This SNP differentially binds the transcription factor, NUDR/Deaf1, leading to cell-type specific effects on transcription in vitro. To investigate the direct effects of this SNP in the heterogeneous cellular context of the brain, we generated humanized transgenic mice using a design that maximized the local transcriptional landscape of the human HTR1A gene while also controlling for effects of genomic insertion location. Expression of the human transgene in a 5-HT1A null mouse resulted in line-dependent expression of human 5-HT1A. The effect of rs6295 on protein levels and behavior similarly differed across lines, suggesting that the penetrance of rs6295 may depend upon background genetic factors. Together, this work confirms that relatively subtle differences in 5-HT1A levels can contribute to differences in behavior and highlights the challenges of modeling human non-coding genetic variation in mice.

## Introduction

Serotonergic systems have been extensively implicated in mood and anxiety disorders, which represent the most common and costly psychiatric disorders. Convergent evidence from humans and animal models suggests that the serotonin 1a receptor plays a primary role in modulating the effect of serotonin on stress and anxiety-related behaviors. Clinically, 5-HT1A agonists, such as buspirone, are prescribed as anxiolytics and as augmentation for antidepressants (Howland, 2015; Trivedi et al., 2006). Likewise, pharamacological and genetic manipulations of the 5-HT1A system in rodents alters anxiety and depression-related behaviors (Heisler et al., 1998; Zhuang et al., 1999). One particularly striking feature of the 5-HT1A system is that relatively small reductions in receptor levels (10 – 40%) in particular brain regions are sufficient to alter behavior (Bortolozzi et al., 2012; Donaldson et al., 2014; Richardson-Jones et al., 2010). For instance, a selective 32% decrease in 5-HT1A autoreceptor levels in the raphe in adulthood, which mimics the natural variation in receptor levels observed in humans (Drevets et al., 2007), increases both resilience to stress and likelihood of SSRI responsiveness (Bortolozzi et al., 2012; Richardson-Jones et al., 2010). Likewise, a ~40% decrease selectively during post-natal development increases later life anxiety-related behavior (Donaldson et al., 2014). This suggests that the 5-HT1A system represents a sensitive substrate in which variation in receptor levels can tune behavior. As such, factors that alter 5-HT1A levels may shape individual differences in behavior and psychiatric disease risk.

One potential genetic factor that has been implicated in modulating human 5-HT1A levels is a common G/C single nucleotide polymorphism (SNP; rs6925), located 1019 basepairs (bp) upstream of the serotonin 1a receptor (HTR1A) translation start site (**Fig. 1A**) (P. R. Albert, 2012). A number of gene association studies have found that the rs6295 G-allele is associated with elevated incidence of suicide attempts and depression and reduce anti-depressant responsiveness, although replication of these effects have not be completely consistent (Donaldson et al., 2016; Drago, Ronchi, & Serretti, 2008; Lemonde et al., 2003; Strobel et al., 2003). Beyond these epidemiological findings, there is evidence that rs6295 is associated with differences in 5-HT1A levels assessed via PET imaging and in post-mortem tissue samples (Donaldson et al., 2016; Kautzky et al., 2017), and in vitro work suggests that it may functionally alter 5-HT1A receptor levels in a cell-type specific fashion through altered binding of transcription factors (P. R. Albert, 2012; Czesak, Lemonde, Peterson, Rogaeva, & Albert, 2006; Philippe et al., 2018). Specifically, the derived G-allele fails to bind the transcription factor, NUDR/Deaf1 (Czesak et al., 2006; Le François, Czesak, Steubl, & Albert, 2008). This leads to greater reporter expression from the G-allele in raphe-derived neurons where NUDR/Deaf1 acts as a repressor, but lower levels in forebrain-derived neurons where it acts as a transcriptional activator (Czesak et al., 2006; Le François et al., 2008). This leads to a model in which the C-allele drives higher levels of 5-HT1A in forebrain neurons but lower levels in raphe neurons (P. R. Albert, 2012), a model that is supported by recent work in post-mortem human tissue (Donaldson et al., 2016).

**Figure 1.**
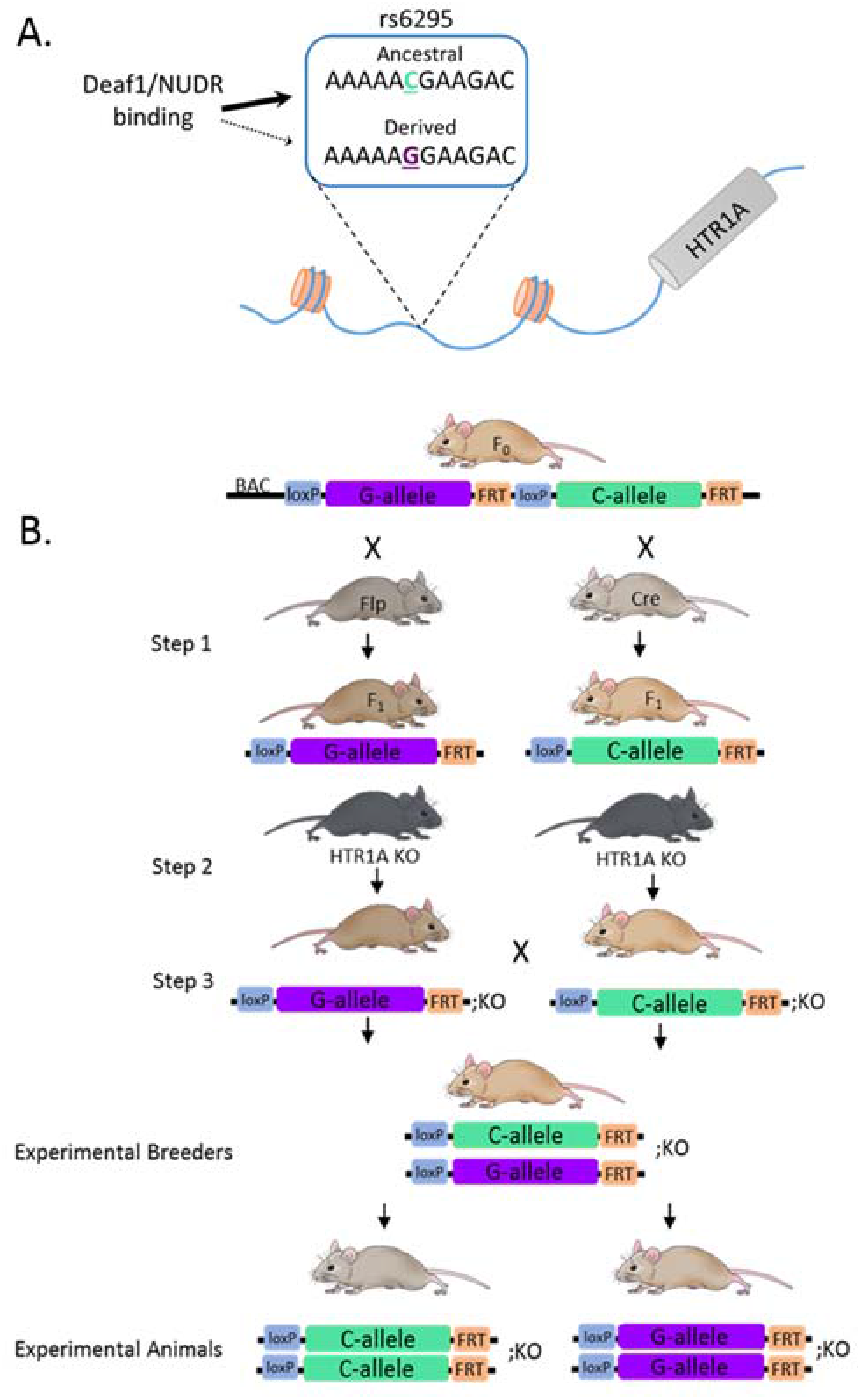
Generation of rs6295 humanized mice. (**A**) The common G/C single nucleotide polymorphism, rs6295, is located upstream of the human HTR1A gene. The transcription factor Deaf1/NUDR binds the C allele but does not bind to the G allele. (**B**) Humanized mice were generated by inserting into the mouse genome a modified bacteria artificial chromosome (BAC) consisting of 180 KB of human DNA containing a duplicated human HTR1A gene containing each allele of rs6295. The rs6295G and rs6295C alleles were isolated by crossing the founder mouse to Cre- and FlpO-recombinase expressing mice (step 1). The offspring of this cross were then bred onto a mouse HTR1A -/- (kO) background (step 2) so that the only source of 5-HT1A in these mice was from human HTR1A locus. Three separate lines were generated.

Despite the strong convergent evidence that rs6295 alters 5-HT1A levels and is associated with psychiatric disease phenotypes, the direct effects of this SNP within the heterogeneous context of the brain remain unknown. To address this, we generated a humanized mouse model of rs6295 by inserting a 180kb bacteria artificial chromosome (BAC) containing the human HTR1A gene into the mouse genome, an approach designed to include relatively distal regulatory elements (**Fig. 1B**). These mice have three important characteristics: 1) the rs6295G and C alleles are present in the same genomic location, 2) we controlled for linked genetic variation in the humanized locus by exclusively varying rs6295, and 3) the only source of 5-HT1A in these animals comes from the human locus. We used this model to investigate the hypothesis that rs6295 directly modulates 5-HT1A levels in a brain-region specific-manner, and that these differences in 5-HT1A levels contribute to differences in stress-coping and anxiety-related behaviors. We found that independent insertion sites led to distinct patterns of 5-HT1A expression in these mice, and that the effects of rs6295 are dependent upon genomic insertion site and/or genetic background. This highlights the complexity of trying to humanize a mouse model to study non-coding variants, but supports the hypothesis that rs6295 can directly modulate 5-HT1A levels and confirms that relatively subtle alterations in 5-HT1A levels contribute to behavior.

## Results and Discussion

### Characterization of h5-HTIA distribution and function in rs6295GC mice

We used a BAC-transgenic strategy to examine the functional effects of rs6295 in the brain. BAC vectors accommodate dispersed regulatory sequences across relatively large regions of the genome (up to hundreds of kb) (Schmidt, Kus, Gong, & Heintz, 2013). This strategy addressed a number of limitations of previous post-mortem human studies and provided the opportunity to examine the effects of rs6295 within the heterogeneous cellular context of the mammalian brain.

5-HT1A is widely expressed in the human and mouse brain, and many of the critical regulatory elements for the expression of HTR1A are well described and conserved between mice and humans (Paul R. Albert & Fiori, 2014). In addition, the general distribution of 5-HT1A receptors is similar between humans and mice, with enrichment in corticolimbic regions, the raphe, and parts of the hypothalamus (Pazos, Cortes, & Palacios, 1985; Pazos, Probst, & Palacios, 1987; Pazos & Palacios, 1985). As such, we anticipated that expression patterns from our BAC would at least partially recapitulate human and mouse expression patterns, acknowledging that patterns of transcription factor expression might differ between species. Specifically, the BAC transgenic approach has previously been used to faithfully recapitulate gene expression patterns across hundreds of mouse genes in the GENSAT project, which reported that screening 3 – 4 founders produces an accurately-expressing line in ~85% of the vectors (Gong et al., 2007). In addition, multiple examples exist of faithful patterns of gene expression have previously been demonstrated for other humanized genetic loci, albeit largely in non-neural tissues(S. M. Lee, Bishop, Goellner, O’Brien, & Pike, 2014). Thus, it was surprising that of the 3 founders generated, they exhibited variable and distinct patterns of 5-HT1A expression, ranging from no adult expression in one line and weak to moderate levels in two other lines. Of the three lines generated, line A and line C contained complete BACs, while the 3’ end of the BAC was truncated in line B (**Supp. Fig. 1**).

We employed hHTR1A, rs6295XX, and h5-HT1A and to indicate the human HTR1A gene, rs6295 genotype, and protein, respectively, and mHTR1A/m5-HT1A to indicate the mouse gene/protein. rs6295GC is the most common genotype in human populations, so we used this genotype to compare our humanized lines (rs6295GC; mHTR1A-/-) to mHTR1A +/- and mHTR1A -/- mice. We mapped the location of 5-HT1A receptors in each line via I^125^-MPPI autoradiography. This revealed distinct patterns of h5-HT1A in each of our lines (**Fig. 2**). Line B exhibited very low levels of h5-HT1A that were limited to regions of the hippocampus and claustrum, while line A had higher levels of h5HT1A that were observable in the hippocampus, amygdala, and prefrontal cortex. Line C was indistinguishable from mHTR1A -/- animals. Due to this lack of h5-HT1A expression, we did not proceed with further analysis of this line.

**Figure 2.**
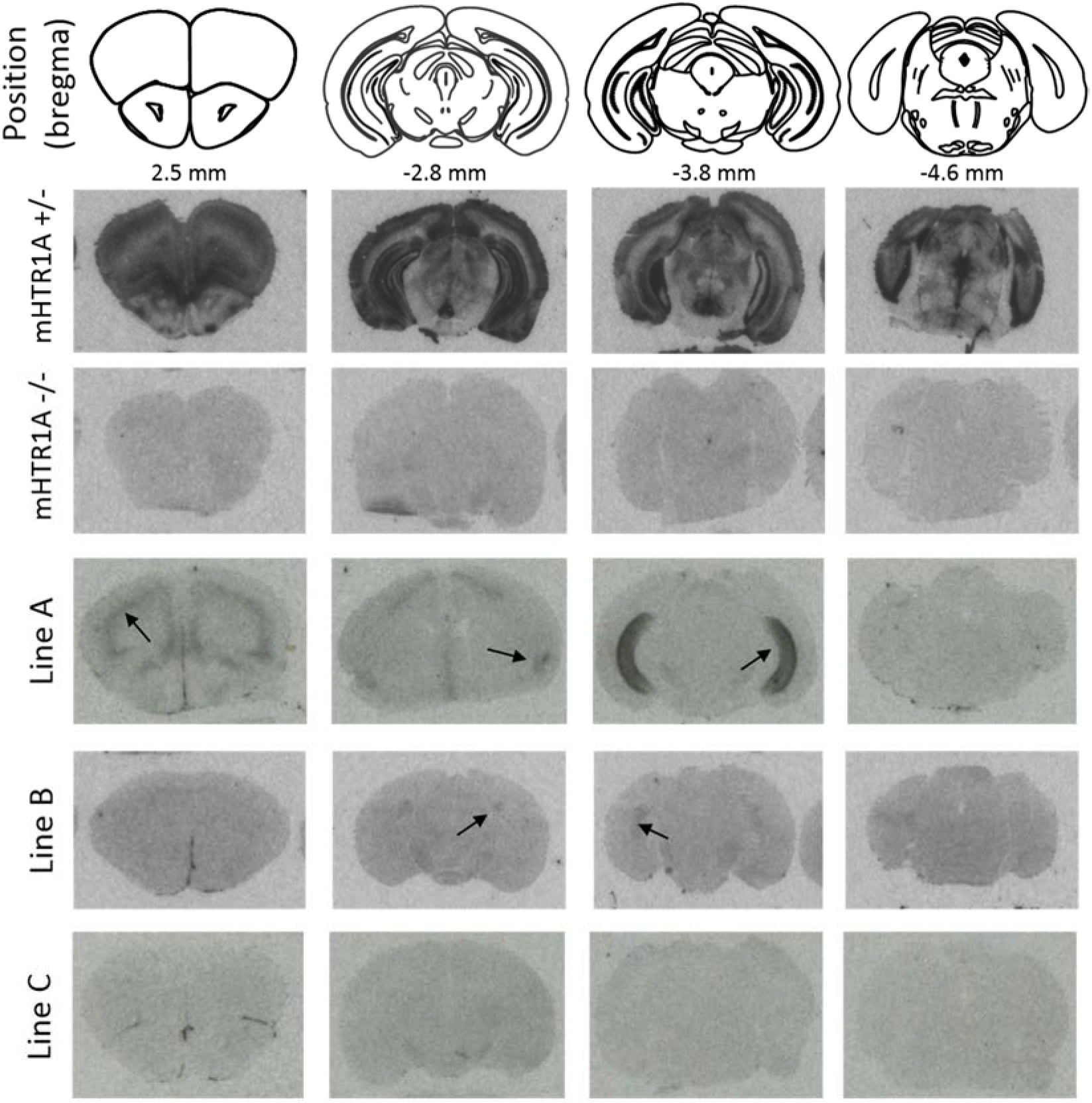
Human 5-HT1A expression varies across transgenic lines. Autoradiography revealed different expression patterns across the three lines created, most likely due to insertion effects of the BAC. Representative images from each line are shown, as well as from mHTR1A -/+ and mHTR1a -/mice. Arrows indicate visually detectable h5-HT1A expression.

Notably, none of our mouse lines expressed h5-HT1A autoreceptors, a result we confirmed using 8-OH-DPAT induced hypothermia (Martin, Phillips, Hearson, Prow, & Heal, 1992). Because expression from the hHTR1A locus was considerably lower than from the mHTR1A locus, we compared our humanized animals to mHTR1A +/- animals, enabling us to use a higher dose of 8-OH-DPAT and maximizing the likelihood of identifying an effect. We observed 8-OH-DPAT induced hypothermia in m5HT1A+/- mice, but hHTR1A; mHTR1A-/- mice were indistinguishable from mHTR1A-/- mice for both line A (m5HT1A+/- = 13; hHTR1A;mHTR1A -/- = 10; m5HT1A-/- = 6; main effect of genotype: F_(1, 26)_ = 41.56; p < 0.001) and line B (m5HT1A+/- = 8 rs6295 = hHTR1A; mHTR1A -/- = 18main effect of genotype: F_(1, 29)_ = 30.561; p < 0.001) lines. (**Supp. Fig. 1**). Full details from statistical analyses are available in **Supplemental table 4**.

### h5-HT1A levels in rs6295CC and rs6295GG mice

Subsequent analyses performed with these lines were limited to h5-HT1A heteroreceptors. We used I^125^-MPPI autoradiography to investigate potential differences in h5-HT1A levels in rs6295CC and rs6295GG mice. Because small changes in m5-HT1A levels during development are known to have long lasting effects into adulthood, we examined levels at two developmental time points, P21 and P60-75 (Donaldson et al., 2014). Line A and B both exhibited h5-HT1A in the ventral hippocampus but differed in detectable receptor expression in the claustrum, cortex, and amygdala. In addition, relative levels of expression differed dramatically between the lines, with receptor levels in line B approximately 1/10^th^ of those observed in line A (**Fig 3**). When we compared GG and CC animals within each line, we found that line B exhibited rs6295-dependent differences in h5-HT1A while line A did not (**Fig 3**). Specifically, line B adolescent rs6295GG animals had higher h5-HT1A protein levels compared to rs6295CC animals in the dorsal hippocampus/subiculum (F_(1, 12)_ = 22.192, p = 0.001) and claustrum (F_(1, 8)_ = 35.472, p = 0.001). In adulthood, line B rs6295GG animals had higher h5-HT1A receptor levels in the dorsal (F_(1, 11)_ = 22.192, p = 0.001), and ventral hippocampus (F_(1, 10)_ = 6.398, p = 0.035), and the claustrum (F_(1, 6)_ = 25.119, p = 0.007). Because these differences were evident during adolescence and adulthood, it suggests that they may be relatively stable across development. **hHTR1A mRNA expression**

**Figure 3.**
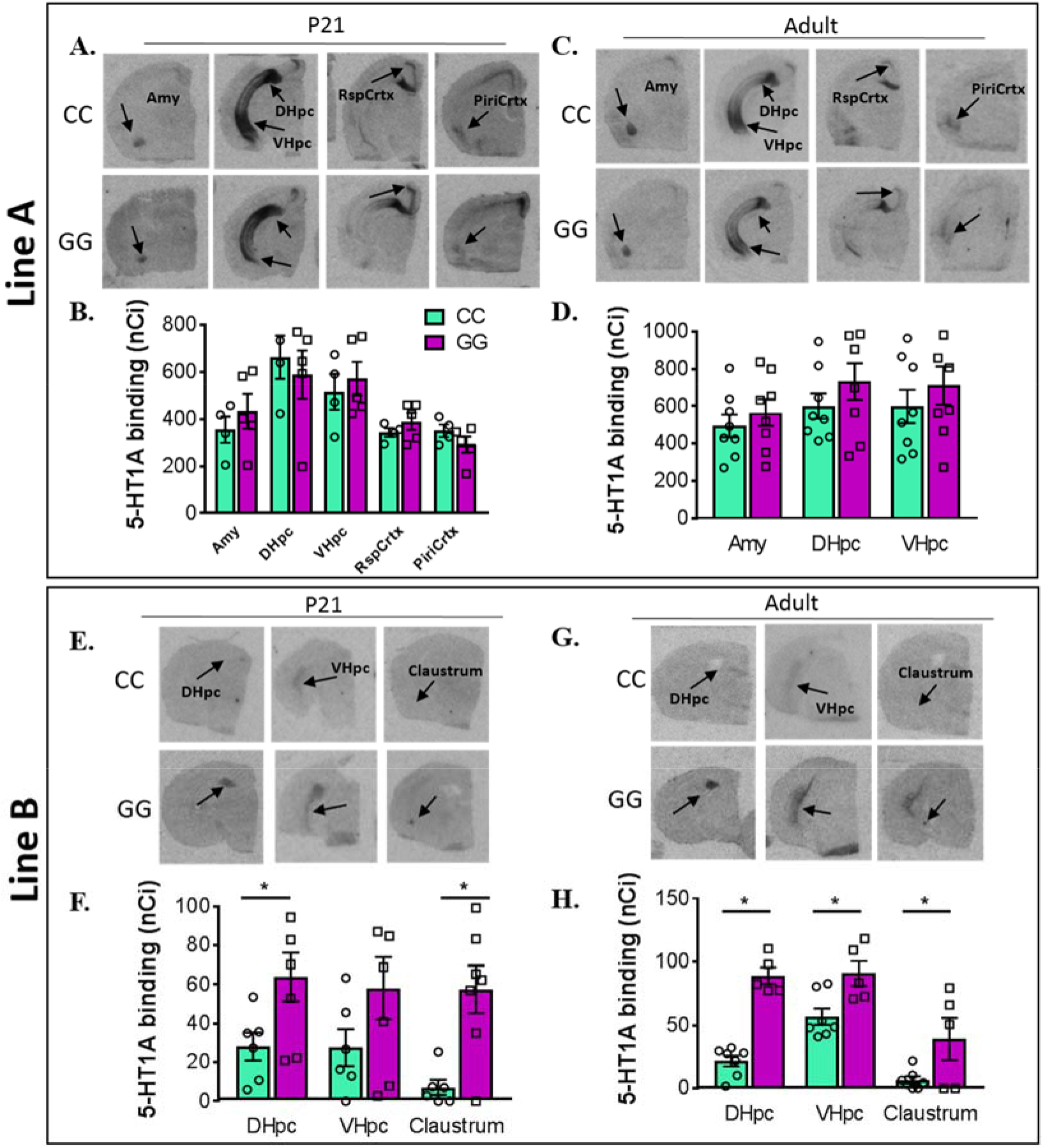
Differences in receptor density between genotypes is line specific. **(A-D)** Autoradiography images (A, C) and quantification (B, D) of h5-HT1A expression in rs6295 GG and CC animals from line A at P21 (A, B) and in adulthood (C, D). No significant differences in h5-HT1A levels were detected in any brain region at either time point. **(E-H)** Autoradiography images (E, G) and quantification (F, H) of h5-HT1A expression in rs6295 GG and CC animals from line B at P21 (E, F) and in adulthood (G, H). At P21, GG mice have higher levels of h5-HT1A in the dorsal hippocampus (DHpc) (p= 0.001) and the daustrum (p= 0.001), but no difference in the ventral hippocampus (VHpc) (p= 0.5290). In adulthood, GG animals have higher h5-HT1A expression in the DHpc (p= 0.0001), VHpc (p= 0.0350), and daustrum (p= 0.0070). Group sizes and statistical analyses available in **supplemental table 4**.

To explore if the differences observed in 5-HT1A receptor protein are accompanied by differences in hHTR1A mRNA, we used quantitative-PCR (qPCR) to examine the expression of hHTR1A using mGAPDH as a control gene. We also examined expression of mDeaf1 as potential confound, reasoning that differences in mDeaf1 levels, as an important transcription factor involved in HTR1A expression, could contribute to differences in hHTR1A mRNA. We did not observe any main or interacting effects of genotype (details available in **supplementary table 4**). In line A, the only significant effect observed was a main effect of brain region on hHTR1A mRNA at p21 (p < 0.0001). In line B, there was a main effect of brain region at both time points (p < 0.05) and a main effect of sex at P21 (p = 0.009) on hHTR1A at P21 (**Supp. Fig. 2**). Deaf1 mRNA also differed as a function of brain region (p < 0.05, Line A P21 and P60, Line B P21 only), but we did not observe any effect of genotype, sex, or their interaction.

While it was surprising that differences in detectable receptor protein levels were not reflected at the level of the mRNA, it is worth noting that levels of HTR1A mRNA in these samples are quite low – often amplifying with a C_t_ >30, which may result in less accurate quantitation. In addition, although hHTR1A mRNA is detectable in the PFC of line B via qPCR, the receptor protein is not detectable above background in this area when using autoradiography. This could be explained by trafficking of the receptor protein to terminals in another brain region or by a failure to insert into the membrane Regardless, it suggests that there is not a direct correspondence between mRNA and protein levels and that differences in mRNA levels in a subset of rs6295-sensitive cells are not sufficient to result in detectable mRNA differences but do contribute to differences at the protein level.

### rs6295-mediated behavioral differences

As the receptor protein is a better indicator of putative functional differences than the mRNA, we examined whether rs6295-mediated differences in h5-HT1A corresponded with differences in anxiety or stress-coping, as detailed below. We found that, mirroring receptor levels as detected by autoradiography, there were no genotype-dependent differences in line A, but line B exhibited subtle differences in anxiety-related and locomotor behaviors. In particular, line B GG animals reared less in the open field and displayed decreased locomotion, a phenotype that has also been observed following manipulations of mHTR1a (Donaldson et al., 2014; Heisler et al., 1998; Richardson-Jones et al., 2010; Zhuang et al., 1999). No genotype-dependent differences were observed in either line for depression-related tests. Together, these data indicate a continuity – in instances in which we observed SNP-dependent differences in h5-HT1A, there is a corresponding difference in behavior, despite extremely low levels of receptor expression. This further confirms the sensitivity of 5-HT1A-mediated behaviors to small differences in receptor levels. Below we report F-statistics and p-value for main effect of genotype; full results of ANOVAs are available in **Supplementary Table 4**.

#### Open Field

rs6295CC animals reared more that rs6295GG animals only in Line B (**Fig. 4**; Line A: F(1,37) = 1.079, p = 0.306; Line B: F(1,37) = 6.386, p = 0.016). No differences were evident in the percent distance traveled in the center (Line A: F(1, 38) = 0.695, p = 0.41; Line B: F(1, 38) = 0.112, p = 0.74), the time in center (Line A: F(1, 36) = 0.093, p = 0.762; Line B: F(1, 38) = 3.643, p = 0.064), or the total distance traveled (Line A: F(1, 38) = 0.655, p = 0.423; Line B: F(1, 38) = 3.643, p = 0.064). As previously observed, locomotion decreased with time (Line A: F(5, 195) = 38.46, p < 0.00001; Line B: F(5, 180) = 114.4, p = < 0.00001), independent of genotype (Line A: F(1, 39) = 0.004, p = 0.951; Line B: F(1, 36) = 1.761, p = 0.193).

**Figure 4.**
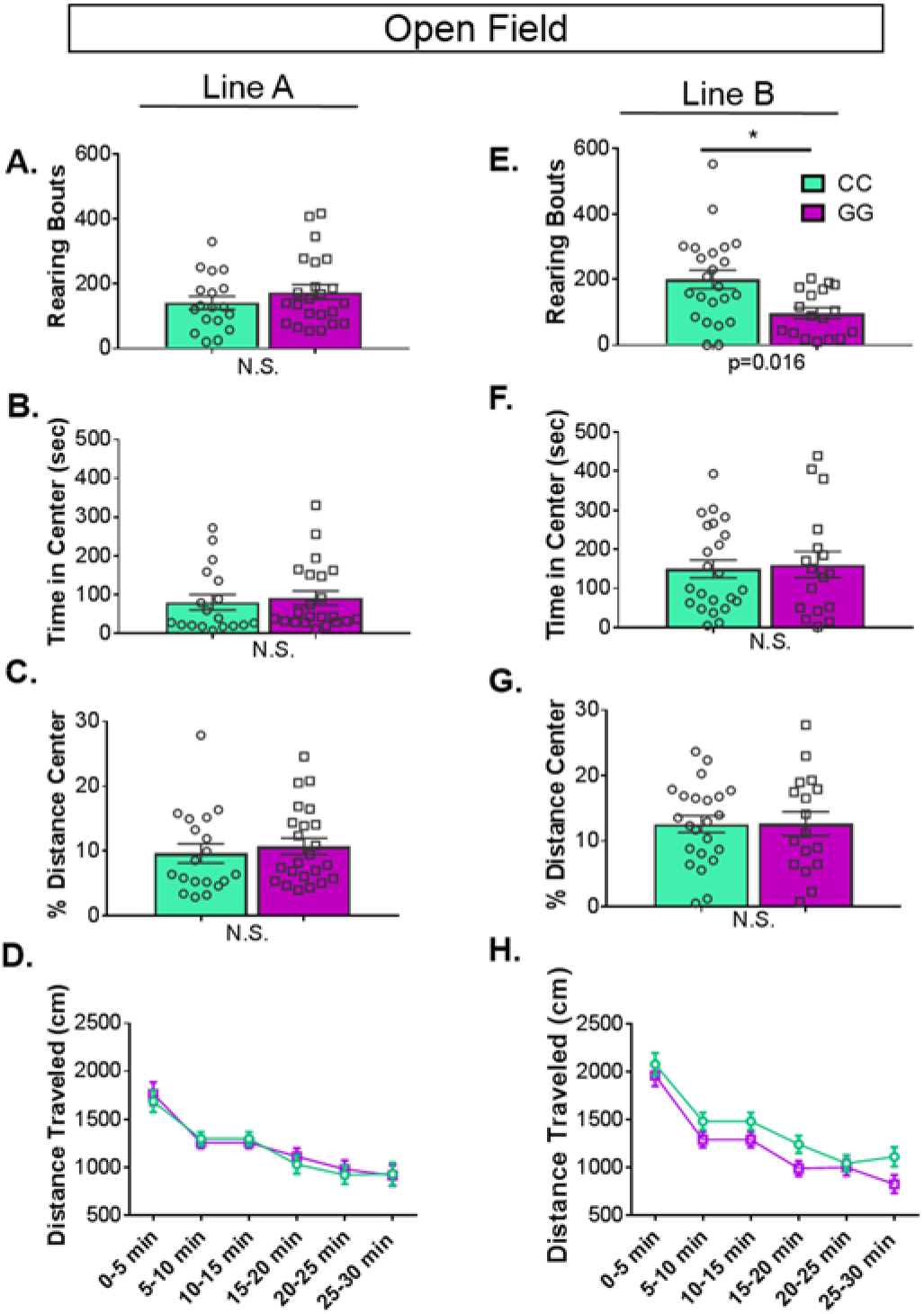
rs6295-dependent differences in rearing in open field are line specific. In open field, the Line A had no statistically significant difference in (**A**) rearing (**B**) time spent in the center, or (**C**) percentage distance traveled in the center between genotypes. (**D**) Likewise, there was no main or interacting effect of genotype on distance travelled over time.(**E**) In line B, CC animals reared more than GG animals (p = 0.016). There were no differences in other measures of anxiety, specifically (**F**) time spent in the center, or (**G**) percentage distance traveled in the center between genotypes. (**H**) There was no main or interacting effect of genotype on distance travelled over time (group sizes and statistics available in **supplemental table 4**). Values represent mean ±SEM.

#### Light-Dark Box

Two animals in line B failed to enter the dark chamber and were omitted from analysis for failure to complete the test. Because data from line B violated the assumption of equal variances, we used a regression to test if genotype significantly predicted behavior. The results of the regression indicate that two predictors explained 22.5% of the variance (R^2^ = .22, F(2, 35) = 5.077, p = 0.012). Genotype significantly predicted total distance traveled (CC>GG, β = −0.445, p = 0.011) (**Fig. 5**). Genotype did not significantly predict percent distance traveled in light (β = −0.192, p = 0.308), light zone entries (β = −0.216, p = 0.197), or total time in the light chamber (β = −0.141, p = 0.455). We analyzed line A using an ANOVA. There were no genotype differences for total distance traveled (F(1, 38) = 0.011, p = 0.916), percent distance traveled in light (F(1, 38) = 0.113, p = 0.739), light zone entries (F(1, 37) = 1.08, p = 0.306), or total time in the light chamber (F(1, 38) = 0.098, p = 0.755)

**Figure 5.**
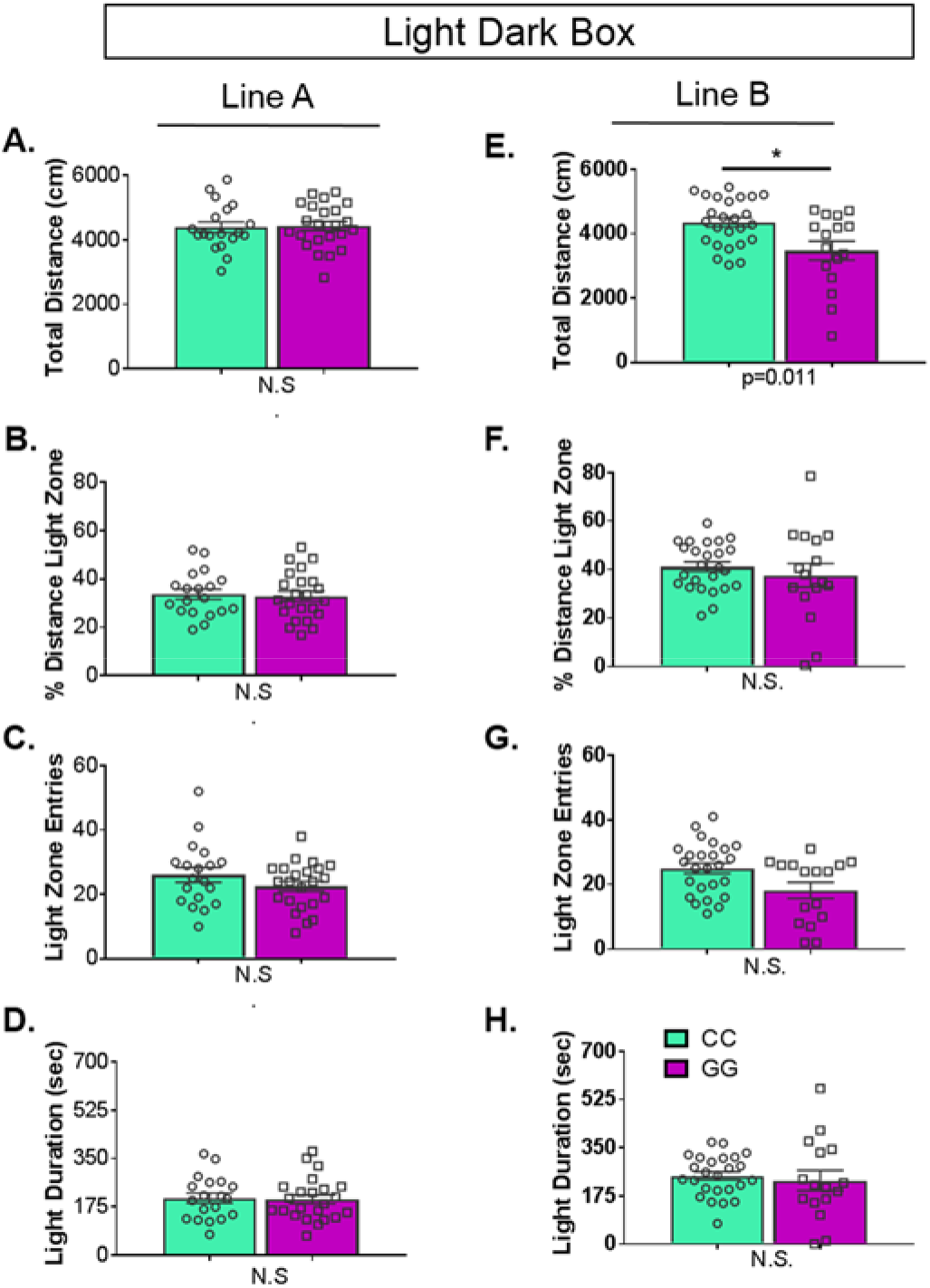
Genotype-dependent differences in light dark box behaviors is line specific. **(A – D)** In Line A, there were no differences between genotypes in any of the metrics examined. In contrast, in line B rs6295CC animals traveled (p = 0.011) There was no difference in **(G)** light zone entries, **(F)** the percent distance traveled in the light or **(H)** light zone duration between genotypes. Values represent mean ±SEM. Group sizes and statistics are available in **supplemental table 4**.

#### Zero Maze

There were no genotype-dependent differences in performance on the elevated plus maze (**Supp. Fig. 4**). Specifically, rs6295CC and GG animals did not differ in the percent time in the open area (Line A: F(1,13) = 0.0.007, p = 0.935; Line B: F(1, 38) = 0.070, p = 0.7920) or the number of entries into the open areas (Line A: F(1,13) = 2.682, p = 0.125; Line B: F(1,37) = 3.643, p = 0.560).

#### Novelty Suppressed Feeding

There was no difference in rs6295CC and rs6295GG mice in the latency to eat food in the novelty suppressed feeding test (**Supp. Fig. 5**). Line A: F(1, 39) = 0.15, p = 0.701; Line B: F(1, 39) = 3.711, p = 0.061. Likewise there was no main effect of genotype for latency to eat in the home cage (Line A: F(1, 35) = 1.112, p = 0.299; Line B: F(1, 35) = 0.026, p = 0.872), the amount eaten (Line A: F(1, 36) = 0.517, p = .0477; Line B: F(1, 39) = 1.75, p = 0.194), or weight loss (Line A: F(1, 38) = 0.655, p = 0.423; Line B: F(1, 38) = 3.643, p = 0.064).

#### Forced Swim Test

Forced swim test was performed on two consecutive days (**Supp. Fig. 6**). As has been previously observed, there was a significant main effect of time on float duration on day 1, with increased floating later in the test for both lines (line A: F(1.76, 54.4) = 37.044, p < 0.0001, line B: F(3.543,131.05) = 11.9; p < 0.0001). However, there was no genotype-dependent differences in floating in forced swim test on either line. (Line A; day 1: F(1, 36) = 0.348, p = 0.559) and day 2: F(1, 36) = 0.207, p = 0.652; Line B: day 1: F(1, 39) = 0.000144, p = 0.99; day 2: F(1, 39) = 0.146, p = 0.705).

#### Tail Suspension Test

There was no difference in time spent immobile between rs6295CC and rs6295GG mice in tail suspension test (**Supp. Fig. 7**). In line A, there was no difference in immobile time between genotypes (F_(1, 36)_ = 0.548; p = 0.464). Similar results were observed in line B; main effect of genotype: F_(1, 39)_ = 1.861; p = 0.18).

Although the direction of our behavioral effects are consistent with epidemiological evidence suggesting that the rs6295G-allele is a risk allele, the relationship between our observed 5-HT1A differences and behavior are not immediately evident. We do not know which cell types are expressing 5-HT1A in our transgenic mice, but the ventral hippocampus has been broadly implicated in modulating anxiety (Jimenez et al., 2018; Kheirbek et al., 2013), and depending upon the cell type expressing 5-HT1A, differences in receptor levels could modulate anxiety by altering hippocampal activity. Alternatively, hippocampal 5-HT1A signaling during early post-natal development is required for normal synaptogenesis, and differences in anxiety could reflect long term impacts of genotype-dependent differences in hippocampal development (Mogha, Guariglia, Debata, Wen, & Banerjee, 2012).

One unanticipated result of this study is that rs6295 modulates h5-HT1A and behavior in only one of our lines. There are two potential explanations for this result. First, each line presumably has a different BAC insertion site, leading to different patterns of receptor expression. It is possible that insertion effects also account for differences in whether rs6295-sensitive transcription factors are being employed during hHTR1A transcription. For instance, if transcription is Deaf1 independent in line A but not line B because of other factors that are part of the insertion-dependent transcriptional complex, that would lead to rs6295-mediated differences in only one line. Alternatively, we controlled for background genetic effects *within* our lines by generating CC and GG mice from heterozygotes. However, given the number of breeding steps needed to generate our lines, it is possible that background differences *between* our lines may contribute to differences in the penetrance of rs6295. Background genetic effects have been shown to moderate the penetrance of other genetic variants. For example, a serotonin transporter (SERT) Ala56 genetic variant that is overtransmitted in autism, causes multiple biochemical, physiological, and behavioral changes in mice with a 129S6/S4 genetic background (Veenstra-VanderWeele et al., 2012), but these phenotypes are not evident when the same allele is bred onto a C57Bl/6 background (Kerr et al., 2013). While we cannot distinguish between these possibilities, we did not observe any differences in Deaf1 expression, so SNP and line effects are not due to differences in Deaf1 transcription. This highlights the challenges related to modeling human cis-regulatory variation in a mouse model where epistatic effects, differences in transcription factor environment, and a lack of direct sequence homology should be considered.

The lack of concordance between h5-HT1A patterns in our humanized mice and those observed in humans and wildtype mice indicates that the model may have limited utility beyond informing the question of whether rs6295 can impact h5-HT1A levels. Thus, in other work, we also explored whether an analogous SNP could be introduced into the native mHTR1A promoter region (Philippe et al., 2018). There are two putative Deaf1 binding sites in the mHTR1A locus, and previous work suggests one of them is functional. The sites do not exhibit direct homology with the human locus, and as a result, there is not a clear single nucleotide change that would model rs6295. Instead, we tested a few different putative mutants, comparing their effects on expression with the rs6295G and C-allele. As detailed in Phillipe et al. (2018) none of our mutations resulted in consistent replication of the effects of rs6295 G-allele across different cell types in vitro. Thus mutating the endogenous mouse Deaf1 binding site(s) as a model for rs6295 would have significant limitations.

In sum, our approach had a number of advantages, and modified versions of our dual Cre-Flp strategy are an innovative way to ensure that two variants and/or haplotypes can be directly compared. As used here, this approach highlights the complexity of modeling human cis-regulatory differences in mice, an important consideration as the majority of GWAS “hits” fall in non-coding genomic regions. Importantly, our results suggest a strong need to investigate potential gene variant effects within an appropriate genomic context. Despite this, our data do support a role for rs6295 in directly modulating receptor levels and further support the finding that subtle differences in 5-HT1A levels can be behaviorally meaningful, making the 5-HT1A system an ideal system for investigating mechanisms that contribute to individual differences in receptor levels in order to further our understanding of psychiatric disease risk.

## Materials and Methods

### Generation of rs6295 humanized mice

We generated transgenic humanized mice carrying a genomically integrated BAC (bacterial artificial chromosome) of ~180 kb containing a duplicated version of the human HTR1A gene flanked by FRT and loxP sites. BAC # RP11-158J3 (BACPAC Resources) was selected as it had the no other protein-coding genes and HTR1A is located in the middle. It was electroporated into SW102 cells for subsequent recombineering (National Cancer Institute: Biological Resources Branch) (E. C. Lee et al., 2001; Yu et al., 2000). There is a loxP site in the RP11 backbone, so we used p23loxZeo (gift from Dr. Kenji Tanaka) to replace the loxP site with a zeocin resistance cassette. Next, we replaced the HTR1A gene-containing-region (6054 bp) with a *rpsL+-kana* marker (Addgene #20871) containing a positive kanamycin resistance marker and a negative streptomycin-sensitive marker (Wang, Zhao, Leiby, & Zhu, 2009). The 5’ (ATG −1944 bp) and 3’ (ATG +4110 bp) ends of this region were chosen because they were the closest regions with low homology outside of known regulatory regions and the 3’ UTR. In parallel, a plasmid containing a duplicated version of the same HTR1A-containing region was also generated. The initial rs6295G allele flanked by a *SalI*-loxP site (5’ end) and an FRT-Xho1 site (3’ end) with 100 bp flanking homology domains was generated via gene synthesis and cloned into pUCminusMCS (pUCrs6295G: Blue Heron Biotechnology). This plasmid was modified via targeted mutagenesis to alter the G-allele to rs6295C, resulting in identical sequences except for rs6295. pUCrs6295G was linearized via *Sal1* and the rs6295C-allele was removed from its backbone via *Sal1* and *Xho1* digestion. This was then ligated together to produce pUCrs6295C:rs6295G, which was then recombined into the BAC, replacing the *rpsL+-kana* marker. Integrity of *HTR1A*, loxP, and FRT sites were confirmed via Sanger sequencing.

Three BAC-transgenic founder mice were generated at the Duke Neurotransgenic Facility and shipped to Columbia University. Upon receipt, founders and their offspring underwent successive breeding. Rates of transgene inheritance in F1 animals are shown in **supplementary table 1**. The founder mice were crossed sequentially to Cre (B6.Cg-Edil5^Tg(Sox2-cre)1Amc^/J; JAX #008454) and FlpO (B6.Cg-TG(Pgk1-flpo)10Sykr/J; JAX # 011065) recombinase-expressing mice, which resulted in F1 generation mice who had one copy of the rs6295G allele or one copy of the rs6295C allele, respectively. Excision was confirmed via a lack of overlapping peaks in Sanger sequencing reaction and Taqman SNP genotyping (assay # 4351379; ThermoScientific Fisher). F1 mice were bred onto an mHTR1A KO (-/-) generated from 5HT1A flox mice (Samuels et al., 2015). F2 mice were bred back to each other, resulting in F3 rs6295 GC; mHTR1A-/- animals for breeding experimental animals. F3 rs6295GC; mHTR1A-/- animals were bred together to generate rs6295GG and CC experimental mice (**Figure 1**). Throughout the rest of this manuscript, rs6295GG or CC mice are on a mHTR1A-/- background unless otherwise specifically stated.

### Husbandry

Mice were housed three - five per cage on a 12-hour dark-light cycle (06:00-18:00) at 20-25°C. *Ad libitum* food and water were provided. All behavioral experiments took place during the light cycle. All animal procedures were approved by Columbia University’s and University of Colorado’s Institutional Animal Care and Use Committee.

### Verification of BAC integrity

In order to determine whether the entire BAC integrated into the genome, we developed a series of 19 primer pairs designed to amplify small fragments tiled every ~ 10 kb across the entire BAC (**supplemental table 2**). Each reaction consisted of 1ul primer pair (10 pmol), 0.5 ul DNA, 0.2 ul Taq DNA polymerase, 3.3 ul ddH2O, and 5ul premix B (Epicentre Biotechnologies) with reaction conditions: 95°C for 1 minutes; 35 X (95°C for 15 sec, 59°C anneal for 30 sec, 72°C for1 min). Bands were resolved on a 3% agarose gel and compared with a positive control PCR amplified from purified BAC-only DNA.

### Genotyping

Toes were taken at postnatal day 7 and digested using 97uL DirectPCR Lysis Reagent-Mouse Tail (Viagen Biotech) and 3uL proteinase K (20 mg/uL Gold Bio Technology Inc). In order to identify mHTR1A -/- mice, genotyping was performed using primers 5’- CACTTGGTAGCTGAAGGTCACG, 5’- TAGCTGGAGCCTCTGAGCGC and 5’- GTACCGGGCGAAGCACTGC. Each reaction consisted of 1ul primer pair (10 pmol), 1 ul DNA, 1.5 ul ddH2O, and 5ul Econotaq (Lucigen Corporation) with reaction conditions: 95°C for 1 minutes, 95 for 15 seconds 64 for 15 seconds; 35 X (95°C for 15 sec, 59°C anneal for 30 sec, 72°C for1 min). Bands were resolved on a 3% agarose gel.

rs6295 genotyping was performed using taqman SNP genotyping (assay # <u>4351379</u>; ThermoScientific Fisher) on a Applied Biosystems QuantStudio 3 qPCR machine. Samples were genotyped in duplicate in MicroAmp Fast Optical 96 well Reaction Plate (Applied Bio Systems) in a rection containing 5uL of DNA, 0.3uL probe SNP Genotyping Assay, 3.4uL Nuclease free H2O 4.3uL Taqman Fast master mix (Life Technologies), and 2uL ddH2O per well for a total volume of 15 uL. The cycling conditions were pre-holding stage: 60°C for 30 seconds, holding stage: 95°C for 10 minutes, 40X (95°C for 15 seconds, 60°C for 1 minute), and post holding stage: 60°C for 30 seconds. Results were compared with positive controls for known rs6295CC, rs6295GG, and rs6295 GC samples, as verified by Sanger sequencing.

### Tissue collection and micro-dissection

Experimental mice were sacrificed as adults (post-natal day 60 - 75; P60) via cervical dislocation followed by decapitation. The left hemisphere was excised and flash frozen and stored −80°C. The pre-frontal cortex and the entire hippocampus of the right hemisphere were micro-dissected by hand and stored at −80°C.

### Autoradiography

Complete brains from rs6295GC mice and left hemispheres from rs6295GG and rs6295CC mice were cryosectioned at a thickness of 20μm and thaw mounted onto Superfrost Plus slides (Fisher Scientific). Sections were maintained at –80 °C until processing. Sections were processed for 4-(2’-methopxyphenyl)-1-[2′-(n-2″-pyridinyl)-p-[^125^I]iodobenzamido]ethylpiperazine (^125^I-MPPI) binding as previously described in other studies (Richardson-Jones et al., 2010). All experimental and control brains for a given line were processed and exposed to film as a single batch. Receptor levels were quantified as follows. Films were scanned at 1600 dpi (Epson Perfection V600 Photo Scanner) and binding intensity was gauged using FIJI software by an experimenter blind to genotype (Schindelin et al., 2012). Levels of 5-HT1AR binding were determined by measuring intensity of the region of interest and subtracting a “background” level from a nearby region in the same section which lacked specific binding. Each region of interest was sampled at least 3 times and the average intensity was used. Measurements were only used if the intensity was on the linear portion of the standard curve obtained from an ARC146-F 14C standard (ARC, St Louis, MO).

### RNA extraction and cDNA generation

Micro-dissected hippocampus and pre-frontal cortex were processed for total purified RNA. The protocol outlined for the Norgen Total RNA/gDNA kit (Norgen Biotek #48700, Thorold, ON, Canada) was used to extract RNA from tissue samples with the following modifications: 600 ul rather than 300 ul of lysis buffer was used and the final elution of RNA was performed twice with 25uL of elution solution rather than a single 50ul elution. 2uL of total RNA for each sample was then nanodropped to determine RNA concentration (Agilent Technologies, Santa Clara, CA). All RNA samples were then standardized to 25ng/ul by dilution with RNAse-free water (Norgen). High Capacity cDNA Reverse Transcription kit (Applied Bio Systems) was used to generate cDNA. cDNA for each sample was produced using a mixture of 2uL 10xTR buffer, 0.8uL dNTP, 2uL random primers, 1uL Transcriptase, 1uL RNA inhibitor, and 3.2uL ddH2O. 250 ng of RNA was then added and cDNA was generated using reaction conditions: 25°C for 10 min, 2x(37°C for 120 min, 85°C for 5 min). Samples were stored at −20°C until processing.

### Quantitative RT-PCR (qRT-PCR)

We examined gene expression in rs6295GG and CC mice. Human HRT1A (IDT; Hs.PT.58.50475698.g), mouse Deaf 1(IDT;Mm.PT. 58.29654925), and mouse GAPDH (Taqman Mm99999915-g1, cat # 4331182) were quantified using qPCR. Efficiencies and primer sequences, where available, are provided in **Supplemental Table 3**. Samples and probes were processed in quadruplicate in MicroAmp Fast Optical 96 well Reaction Plate (Applied Bio Systems) with 0.25uL of cDNA, 0.25uL probe, 2.5uL Taqman Fast master mix (Life Technologies), and 2uL ddH2O per well for a total volume of 5uL. The plate was covered with Optical Adhesive Film (Applied Bio Systems), vortexed and centrifuged prior to PCR amplification in an Applied Biosystems QuantStudio 3 qPCR machine (Applied Bio Systems). DNA was amplified in a qPCR reaction with and without addition of cDNA to ensure there was no contamination from genomic DNA. Cycling conditions were as follows: 50°C for 2 min; 95°C for 20s 40 cycles of 95°C for 1s and 60°C for 20s.

The mean cycle threshold (C_t_) value was taken for each sample in quadruplicate. Quadruplicates that had a standard deviation greater than 0.30 were checked for an obvious outlier, and if none could be visually identified, the sample was excluded from analysis. 2^ΔΔC_t_ method was used for analysis, with the hippocampus of CC individuals used as the control group. A two-way RM-ANOVA was used to examine the effects of genotype and sex with brain region as a repeated measure.

### 8-OH-DPAT induced hypothermia

To examine whether our humanized lines expressed functional 5-HT1A autoreceptors, we compared 8-OH-DPAT hypothermic responses in animals with one endogenous copy of the mouse allele, one copy of the human allele on an m5-HT1A -/- background, and m5HT1A -/- mice. 8-OH-DPAT induced hypothermia is mediated exclusively by 5-HT1A autoreceptors (Martin et al., 1992). Body temperature was measured rectally three times to establish a baseline temperature. 5mg/kg of 8-OH-DPAT (Sigma) was administered via intraperitoneal (IP) injection. Temperature was then measured every 10 minutes for 60 minutes. Change in temperature relative to baseline was compared using a RM-ANOVA with genotype and sex as the between subject factors and time as the within subject factor.

### Behavioral phenotyping

Adult mice (postnatal day 60 - 75) underwent a battery of depression and anxiety-related behavior tests. Prior to testing, female mice were condensed into the same cage; male mice were housed with the same animals they were weaned with to avoid eliciting aggression. The tests were administered in the following order: open field under low light, light dark box, elevated zero maze, novelty suppressed feeding, forced swim test, and tail suspension test. All testing apparatuses were cleaned with disinfectant and allowed to dry in between animals.

#### Open field

Open field behavior was tested in square chambers (16 x 16 in) with metal floors, no bedding, and clear walls (Med Associates, Vermont, USA). Low light conditions consisted of ~30 lux in the center of the open field chamber. On testing day, mice were given 30 minutes to acclimate in the behavior room and lighting conditions. Animals were placed in the center of the box and allowed to freely explore the test chamber for 30 minutes. Locomotion and rearing were recorded via beam-breaks using the software Activity Monitor (Med Associates).

#### Light dark box

Testing was conducted in the open field chambers described above with a dark plastic box insert opaque to visible light but transparent to infrared motion tracking beams in half of the chamber. This was joined via an open doorway to the light side of the chamber, which was brightly lit with a fluorescent bulb (~400 lux). Mice were allowed to acclimate to the behaver room for 30 minutes prior to behavior testing. For testing, mice were placed in the light side of the chamber facing away from the door and were allowed to freely explore for 10 minutes. Locomotion, rearing, time, and entries into each chamber were recorded via beam-breaks using Activity Monitor. Mice were omitted from analysis if they failed to enter the dark chamber.

#### Elevated Zero Maze

An elevated zero maze was designed with Tinkecard software and 3D printed with light gray Makerbot filament at Columbia University Engineering Library on a Makerbot printer with the following dimensions (13.5 in. high, 21.5 in. outer diameter, 2 in. arm width, 6 in. tall closed arm walls, with 1cm ridges lining the edges of the open arms. On test day, mice were given 30 minutes to acclimate in the behavior room before being placed on the open portion of the maze. Mice were allowed to explore the maze for 10 minutes. Behavior was recorded and scored using Noldus Ethovision XT software. Independent samples t-tests were used to determine if there were statistically significant differences between genotypes in open arm entries and percent time spent in the open arms.

#### Novelty suppressed feeding (NSF)

NSF testing took place in a shallow white arena (17.875 in. long, 11.670 in. wide x 7.5 in. deep) with a lux of 1200-1300 in center of the box. 13.5 hours prior to testing, mice were weighed, and food was removed from the home cage. On testing day, mice were weighed prior to testing. Animals were removed from their home cage and transferred to the arena with corn bedding lining the bottom. A single pellet of food was placed on a white paper platform positioned in the center of the crate and animals were able to explore for up to 6 minutes. The time at which the mouse took its first bite of food was recorded, and the food pellet was removed. After six minutes or after the mice took a bit of food, they were transferred back to their home cages and allowed to explore for up to 5 minutes. The time at which the mouse took its first bite of food in the home cage was recorded. After 5 minutes, the food was removed from the cage and weighed to determine the amount eaten. The mouse was weighed and change in weight as a percentage of body weight was calculated for each animal. Independent samples t-tests were used to determine if there were statistically significant differences between genotypes in latency to eat in the home and anxiety cages, food eaten/body mass, and percentage weight change.

#### Forced swim test (FST)

The FST paradigm was comprised of a plastic container 7.5 in. wide and 9 in. deep, filled with 23–25°C water. Mice were placed in the bucket and filmed for 6 minutes. The height of water was selected so that animal was prevented from touching the bottom of the plastic tank and also to prevent its escape. This protocol was repeated 24 hours later as previous research suggests that the second testing session may be more sensitive for detecting antidepressant effects (Borsini, Podhorna, & Marazziti, 2002). Behavior was recorded and scored using an automated Viewpoint Videotrack software package (Montreal, Canada). Floating was examined in one minute bins and analyzed using a two way repeated measures ANOVA with time bin as a within-subject measure and genotype as a between-subject measure.

#### Tail Suspension Test (TST)

Mice were suspended by their tail using tape to secure them to a horizontal bar. The animals were suspended for 6 minutes and immobility during this period was assessed using an automated Viewpoint Videotrack software package.

### Statistical analysis

All statistical analyses were carried out using SPSS 24. Outliers were identified by finding the interquartile range (IQR). IQRx1.5 was then added to quartile 3 (Q3) to get the upper range and IQRx1.5 subtracted from added to quartile1 (Q1) gave the lower range; anything outside of these ranges was excluded as an outlier. Residual normalcy was checked, a Kolmogorov-Smirnov value > 0.200 was considered normal. When performing ANOVAs, if the residuals were not normally distributed, the data was transformed until a normal distribution could be achieved. If transformation was not successful in generating a normal distribution, analysis was done using an *ANOVA* with bootstrapping simple sampling method (5000 repetitions, 95% confidence interval) with genotype and sex as the dependent variables. In all cases where two way *ANOVA* was appropriate, sphericity was assumed if sphericity was p > 0.05. If this requirement was not achieved, Greenhouse-Geisser F and p-values are reported. Likewise, if homogeneity of variance was violated, we instead used a regression. Details are provided in **Supplementary Table 4**.

This is an equation line. Type the equation in the equation editor field, then put the number of the equation in the brackets at right. The equation line is a one-row table, it allows you to both center the equation and have a right-justified reference, as found in most journals.

## Acknowledgments

We would like to thank the animal care staff at New York State Psychiatric Institute and at University of Colorado Boulder. We are also grateful to Meghin Sadsad, Eric Klein, Katelyn Gordon, and Kylia Ahuna for providing additional experimental and analytical support. We also thank Paul Albert for thought discussion of experimental design and interpretation of results and Kenji Tanaka for advice on transgenic design. This work was made possible through the support of American Foundation for Suicide Prevention, R00 MH102352, and R00 MH102352-S1 to ZRD.

